# Nitrogen deposition reshapes plant nutrient acquisition strategies: a meta-analysis within the root economics space

**DOI:** 10.1101/2025.03.06.641811

**Authors:** Hui Guo, Yun Zhao, Bangxiao Zheng, Yin Huang, Xiaoqing Chen, Lei Wang

**Affiliations:** School of Environmental Science and Engineering, Xiamen University of Technology, Xiamen 361024, China; Graduate Program in Environmental Science, College of Environmental Science and Forestry, State University of New York, Syracuse, NY 13210, USA

**Keywords:** nitrogen deposition, fine roots, nutrient acquisition strategies, root traits, root economics space, phosphorus addition

## Abstract

Global nitrogen (N) deposition has fundamentally reshaped plant nutrient acquisition by altering mechanistic trade-offs between root exploration-exploitation strategies and mycorrhizal symbiosis. Through a meta-analysis of 135 studies spanning 153 sites, we demonstrate that N deposition stimulates root exploration (+84.6% root length density) and exploitation (+13.2% nitrogen content and +8.5% root biomass) while suppressing mycorrhizal dependence (-32.8% biomass, -18.73% colonized root length, -7.55% hyphal length), indicating a systemic shift toward root-autonomous nutrient acquisition. Divergent responses emerged between plant lifeforms: woody plants prioritize exploitation (+13.2% nitrogen content) over exploration capacity (-7.9% root length density), whereas herbaceous species exhibit synergistic enhancement of both strategies (+88% root length density, +13.2% nitrogen content and +71.3% root biomass). In contrast to N-only effects, Combined nitrogen and phosphorus additions reversed mycorrhizal suppression (+41.9% biomass), highlighting their persistent role in phosphorus acquisition under elevated nutrient conditions. The root economics space framework demonstrates predictive power, with plants possessing lower specific root length (SRL) and higher nitrogen content exhibiting stronger positive responses to N deposition, while those possessing higher SRL and lower nitrogen content showed reduced investment in root systems. These trait-mediated thresholds in carbon-nutrient tradeoffs refine our capacity to model belowground ecological responses to anthropogenic nitrogen perturbation, establishing a mechanistic basis for projecting ecosystem trajectories under global change.

## 1. Introduction

Plant nutrient acquisition operates through two principal pathways: direct uptake via fine roots and symbiotic associations with mycorrhizal fungi (Chen et al., 2016; Averill et al., 2019). Functionally specialized fine roots execute dual strategies in soil resource acquisition: nutrient “exploration” through architectural expansion and “exploitation” via physiological optimization (Putz et al., 2024). The exploration strategy involves maximizing root system coverage through increased root length density, enabling plants to access spatially heterogeneous nutrients through low-cost root construction. This is typically characterized by morphological adaptations such as reduced root tissue density, smaller diameter, and elevated specific root length (SRL) (Freschet et al., 2021). Conversely, exploitation strategies prioritize biomass investment in nutrient-rich roots, with uptake efficiency strongly correlated to root nitrogen content (Griffiths et al., 2021; Yi et al., 2024). Complementing these root-based mechanisms, mycorrhizal symbioses extend plants’ nutrient foraging capacity through extensive hyphal networks, particularly enhancing phosphorus acquisition (Genre et al., 2020).

Anthropogenic nitrogen (N) deposition has emerged as a critical modifier of terrestrial ecosystems, elevating global N inputs by >50% since pre-industrial times (Galloway et al., 2008). This perturbation fundamentally alters soil biogeochemistry, triggering cascading effects on plant nutrient acquisition strategies. Elevated N availability typically enhances fine root nitrogen content while exerting minimal influence on root diameter, tissue density, and SRL (Li et al., 2015; Zhao et al., 2022). Although fine root biomass shows limited responsiveness to N enrichment (Liu & Greaver, 2010; Peng et al., 2017), accelerated root turnover rates suggest altered carbon allocation patterns (Ma et al., 2021). Notably, N deposition induces substantial declines in mycorrhizal colonization (Han et al., 2020), indicating a strategic shift from fungal-mediated nutrient acquisition towards root-centric strategies. However, the mechanistic interplay between exploration-exploitation trade-offs and mycorrhizal disengagement under N deposition remains poorly resolved.

Divergent evolutionary pressures have shaped contrasting nutrient acquisition strategies between herbaceous and woody plants. Herbaceous species, constrained by ephemeral life cycles, employ rapid nutrient capture through transient root proliferation (Jach-Smith & Jackson, 2018). N enrichment typically amplifies this “opportunistic” strategy through enhanced root length density (Gao et al., 2023). In contrast, woody plants exhibit conservative resource management, relying on persistent root architectures and mycorrhizal networks to maintain nutrient supply continuity (Guo et al., 2024). These phylogenetic differences suggest fundamentally distinct response thresholds to atmospheric N loading.

Emerging evidence highlights phosphorus (P) availability as a critical modulator of N deposition effects. As ecosystems transition from N- to P-limitation under chronic N enrichment (Peñuelas et al., 2013), combined N+P inputs may recalibrate plant-microbial interactions. While some studies report synergistic N+P effects on mycorrhizal dependence (Camenzind et al., 2016), others document reduced root investment under elevated nutrient availability (Phillips & Fahey, 2007). This apparent paradox underscores the need to disentangle stoichiometric interactions in plant nutrient strategies.

The root economics framework provides a predictive lens for interpreting trait-mediated adaptations (Bergmann et al., 2020). Along the nutrient acquisition axis, plants exhibit a continuum from mycorrhizal-dependent strategies (thicker roots, lower SRL) to self-reliant strategies (thinner roots, higher SRL). N deposition likely disadvantages mycorrhizal-dependent species through fungal suppression (Lilleskov et al., 2019), while favoring plants with exploitative root traits. Parallel to leaf economic spectra, root nitrogen content and tissue density reflect “fast-slow” resource use strategies (Wright et al., 2004). Fast-strategy plants with high nitrogen roots may gain competitive advantages under N enrichment (Ordonez & Olff, 2013), though root trait response patterns remain inadequately quantified.

To address these knowledge gaps, we conducted a global meta-analysis of 135 studies examining N deposition effects on nine critical belowground traits. We hypothesize that: (a) N enrichment drives strategic shifts from exploration to exploitation, concomitant with mycorrhizal disengagement; (b) Woody plants optimize uptake via nitrogen enrichment, while herbaceous species enhance both exploration and exploitation; (c) N+P co-addition partially restores mycorrhizal dependence; (d) Root economic space positioning predicts directional responses, with mycorrhizal-aligned species showing negative responses and self-reliant species exhibiting positive responses; (e) Fast-strategy plants demonstrate amplified biomass responses under N loading compared to slow-strategy counterparts.

## 2. Methods

### 2.1 Data collection

We systematically compiled experimental data through comprehensive searches in Web of Science and China National Knowledge Infrastructure (CNKI) databases, encompassing studies published through June 2024. The Boolean search strategy combined nitrogen perturbation terms (“nitrogen fertilization” OR “nitrogen addition” OR “nitrogen application” OR “nitrogen supply” OR “nitrogen input”) with belowground component descriptors (“root” OR “fine root” OR “absorptive root” OR “mycorrhizal fungi” OR “arbuscular mycorrhiza” OR “ectomycorrhiza”). The following criteria were utilized during data screening: (1) Field-scale manipulative experiments excluding greenhouse/pot studies; (2) Minimum experimental duration of 12 months; (3) Focus on N-only or N+P treatments in multi-factor designs; (4) Root diameter specifications (≤2 mm, ≤1 mm, ≤0.5 mm) or absorptive root orders (1, 1-2, 1-3); (5) Quantitative reporting of means with variability measures (SD/SE). For non-tabulated data, we employed Engauge Digitizer 11.1 (Mitchell et al., 2017) for graphical data extraction. The final dataset comprised 135 peer-reviewed studies across 153 globally distributed sites (Fig. S1), categorized into three functional trait groups (Table 1): (1) Fine root traits comprising diameter, specific root length (SRL), root tissue density (RTD), and nitrogen content, which collectively define the root economic space (Bergmann et al., 2020); (2) Root system traits operationalized through fine root biomass and root length density, nutrient exploitation capacity was explicitly defined by integrating biomass and nitrogen content based on their established relationship with uptake kinetics (Griffiths et al., 2021; Han & Zhu, 2021; Yi et al., 2024); (3) Mycorrhizal symbiosis metrics including colonized root length (derived from mycorrhizal colonization rate × RLD), fungal biomass, and hyphal length, quantifying fungal-mediated nutrient acquisition pathways.

**Table 1.** Abbreviations, full names, trait types, and units of belowground plant characteristics.

| Abbreviation | Full name | Traits type | Unit |
| --- | --- | --- | --- |
| RD | root biomass | Root trait | mm |
| SRL | specific root length | Root trait | m g<sup>-1</sup> |
| RTD | root tissue density | Root trait | g cm<sup>-3</sup> |
| RNC | root nitrogen content | Root trait | % |
| RB | root biomass | Root system trait | g m<sup>-2</sup> |
| RLD | root length density | Root system trait | m m<sup>-2</sup> |
| RLCAF | root length colonized by arbuscular mycorrhizal fungi | Mycorrhiza trait | m m<sup>-2</sup> |
| FHLD | fungal hyphal length density | Mycorrhiza trait | cm g<sup>-1</sup> |
| FB | fungal biomass | Mycorrhiza trait | nmol mg<sup>-1</sup> |

These nine trait variables enabled a comprehensive analysis of plant nutrient acquisition strategies under N deposition. A total of 135 studies meeting the above criteria were selected, encompassing 153 sites as shown in Fig. S1.

### 2.2 Meta-Analysis

In this study, a meta-analysis approach was employed to analyze the responses of plant belowground traits to N deposition, utilizing the “metafor” package in R. The analysis involved the following steps:

First, the natural logarithm of the response ratio (LnRR), was used as the effect size to measure the influence of N deposition on plant nutrient acquisition strategies. The formula is:

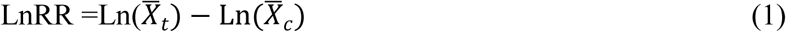

where *X_t_* and *X_c_* represent the mean values for the treatment and control groups, respectively. The variance (*v*) was calculated as:

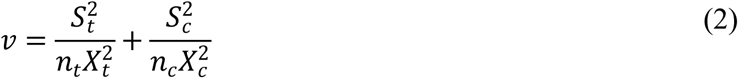

where St and Sc are the standard deviations, and nt and nc are the sample sizes of the treatment and control groups, respectively.

The weight for each observation (wij) was calculated as the inverse of the variance:

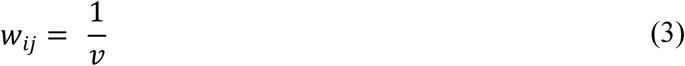

For locations with multiple observations, weights were adjusted by dividing by the total number of observations at each location:

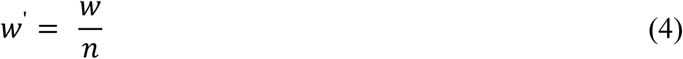

where *n* is the total number of observations at the location. The weighted effect size (lnRR’) was calculated as:

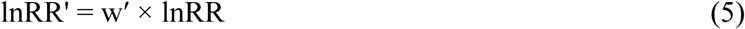

The average effect size (RR++) was computed as:

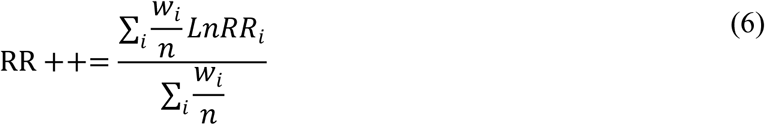

Here, wi and lnRRi are the weights and natural log response ratios for the ith observation.

A random-effects meta-analysis model was employed. When the 95% confidence interval (CI) of an effect size did not include zero, the effect of N deposition on the trait was considered statistically significant. A positive effect size indicates a positive effect, and a negative effect size indicates a negative effect.

To interpret changes in belowground traits under N deposition, we converted the average effect size into percentage change using the formula:

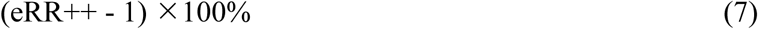

Liner mixed-effects models were employed to evaluate the effects of N deposition amount and duration on belowground traits and can be expressed as:

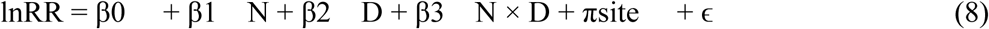

where β represents the coefficients, πsite is the random effect for each site, and ɛ is the sampling error.

Mean imputation was used to caculate any missing standard deviations, sdm:

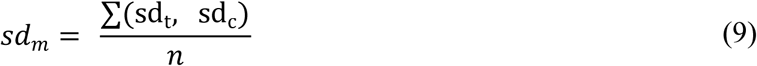

where sdt and sdc are the standard deviations of the treatment and control groups, and n is the number of non-missing values. Missing standard deviations were replaced with sdm.

### 2.3 Data analysis

The dataset was stratified into two experimental cohorts: N-only treatments and N+P co-addition trials. To isolate phosphorus-mediated effects, responses under N+P conditions were systematically benchmarked against N-only treatments through contrast analysis. Subsequent mechanistic investigations focused exclusively on the N deposition dataset.

To elucidate functional differences in nutrient strategy regulation, plants were categorized by growth form: woody species and herbaceous plants. Following the mycorrhizal classification framework of Soudzilovskaia et al. (2020), woody plants were further categorized by symbiotic type: arbuscular mycorrhizal (AM) associates and ectomycorrhizal (EM) symbionts.

Trait response magnitudes were calculated relative to unfertilized control baselines, enabling quantification of N-induced plasticity. A rigorous outlier screening protocol identified Study 25 (Appendix data) as containing statistical outliers (Cook’s distance > 4.7), prompting its exclusion from regression models to maintain analytical integrity.

## 3. Results

### 3.1 Nitrogen deposition reshapes plant nutrient acquisition strategies

Nitrogen (N) deposition enrichment elicited fundamental reorganization of plant nutrient acquisition strategies (Fig. 1). Root exploration capacity, quantified by root length density (RLD), increased by 84.56% through synergistic improvements in specific root length (SRL, +1.52%) and fine root biomass (+8.51%). Concurrently, exploitation capacity intensified with 13.22% elevation in root nitrogen content. This root-centric shift coincided with substantial mycorrhizal disengagement: colonized root length declined by 18.73%, hyphal length by 7.55%, and fungal biomass by 32.76%, indicating a shift toward root-based nutrient acquisition.

**Figure 1.**
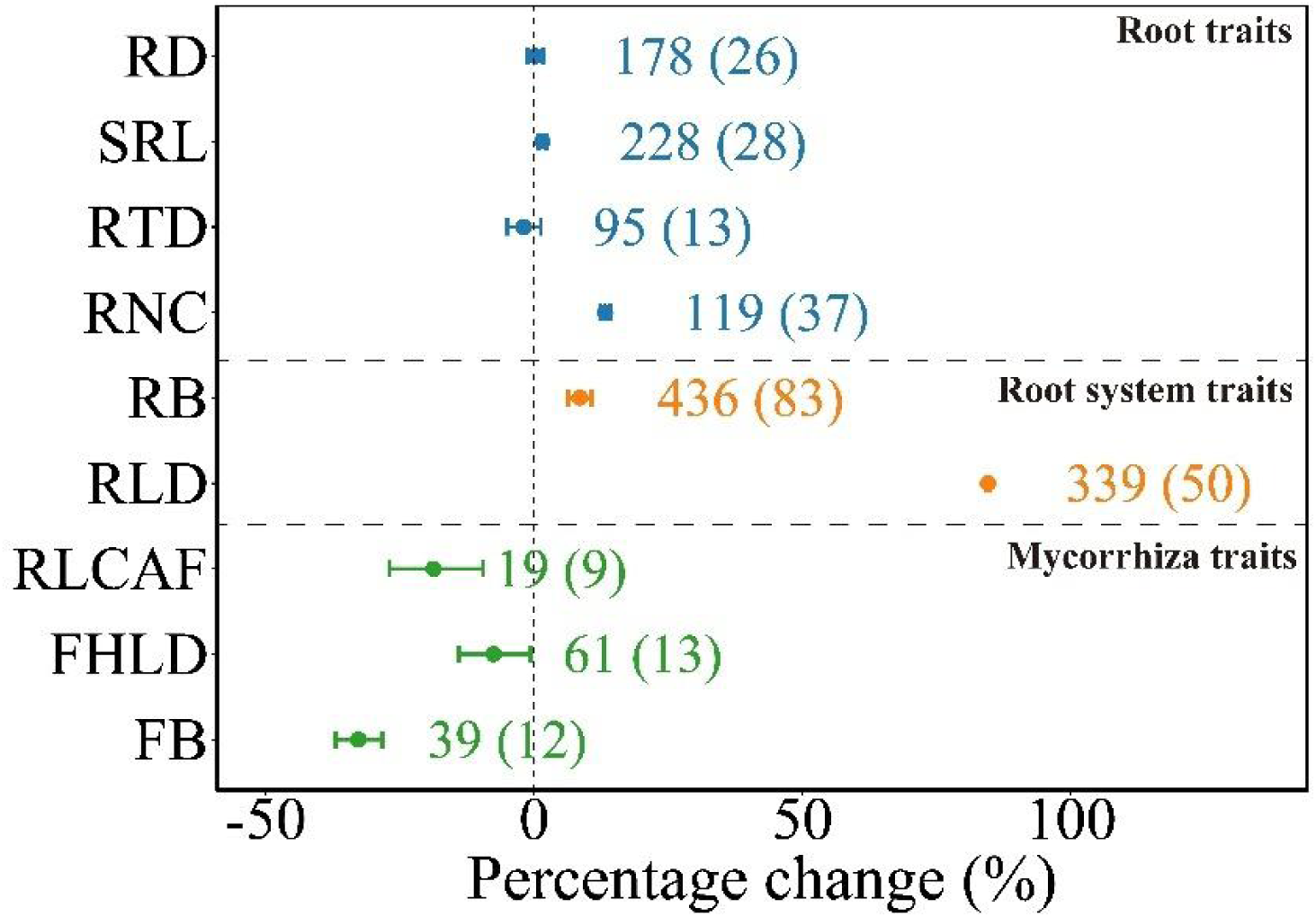
Effect sizes of fine root and mycorrhizal fungal traits in response to nitrogen deposition. Error bars indicate 95% confidence intervals (CIs), with effects considered significant when CIs do not overlap with 0. Numbers within and outside the parentheses represent the sample site and observations for each trait.

### 3.2 Phosphorus co-limitation modulates N deposition effects

Combined nitrogen and phosphorus addition (N+P) raised exploration capacity enhancements (+42.33% RLD), driven by greater SRL (+8.99%) and biomass responses (+19.39%) (Fig. 2). Contrasting with N-only treatments, mycorrhizal biomass increased 41.94% under N+P, suggesting fungal-mediated phosphorus scavenging. In woody plants, N+P induced architectural optimization through 7.17% root tissue density reduction and 8.99% SRL elevation, yielding 25.48% RLD improvement (Fig. S2). This synergistic effect demonstrates phosphorus availability’s critical role in moderating plant-fungal interactions.

**Figure 2.**
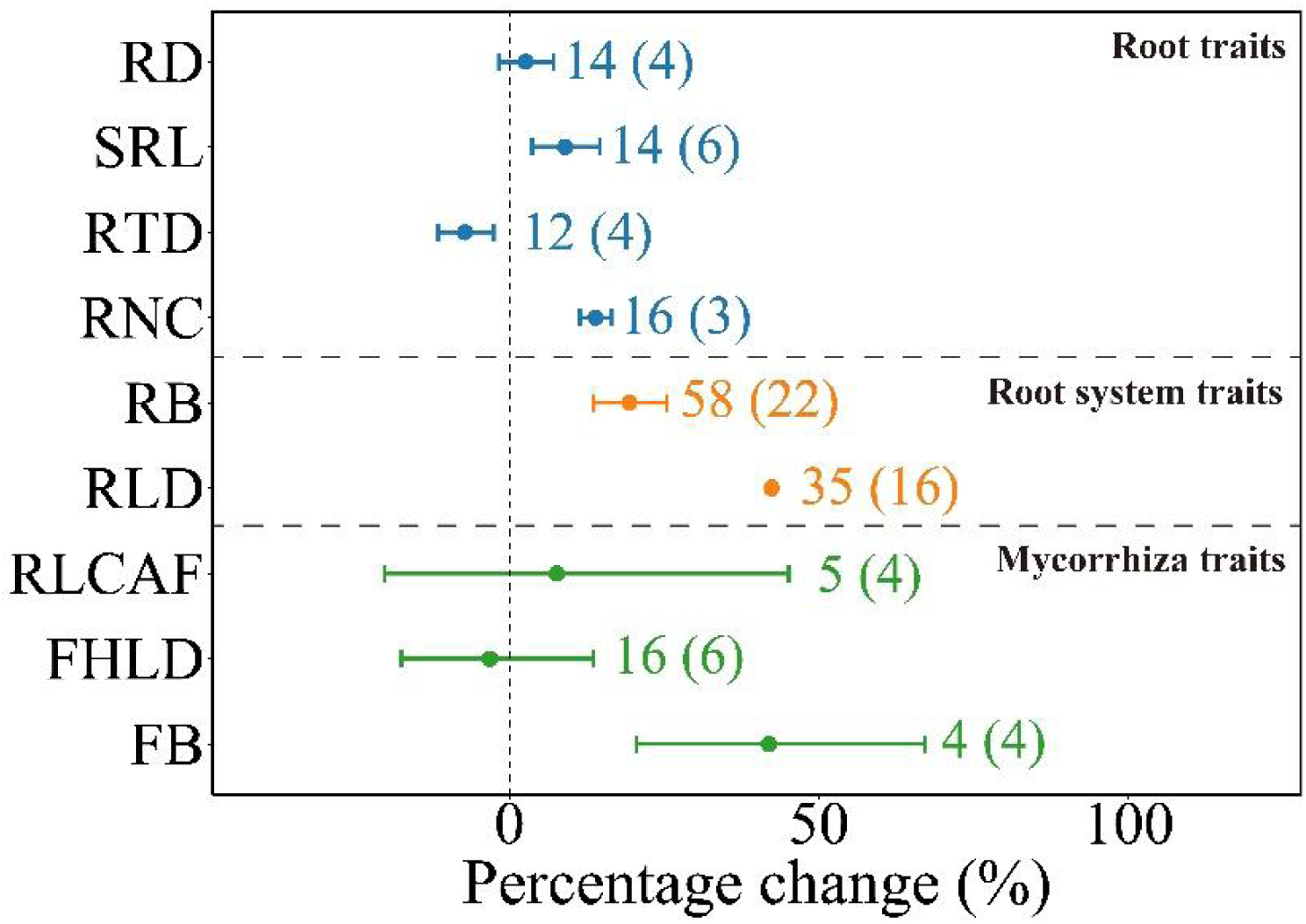
Effect sizes of fine root and mycorrhizal fungal traits in response to combined nitrogen and phosphorus additions. Error bars indicate 95% confidence intervals (CIs), with effects considered significant when CIs do not overlap with 0. Numbers within and outside the parentheses represent the sample site and observations for each trait.

### 3.3 Growth form mediates belowground adaptation strategies

Woody plants exhibited conservative adaptation: Unchanged fine root biomass accompanied by 11.85% tissue density increase, resulting in 7.87% RLD contraction (Fig. 3a). This structural consolidation contrasted with 13.21% nitrogen content elevation, suggesting metabolic intensification rather than spatial expansion. Mycorrhizal dependence declined uniformly (-20.30% biomass), though arbuscular mycorrhizal (AM) species showed unexpected RLD gains versus ectomycorrhizal (EM) counterparts (Fig. S3).

**Figure 3.**
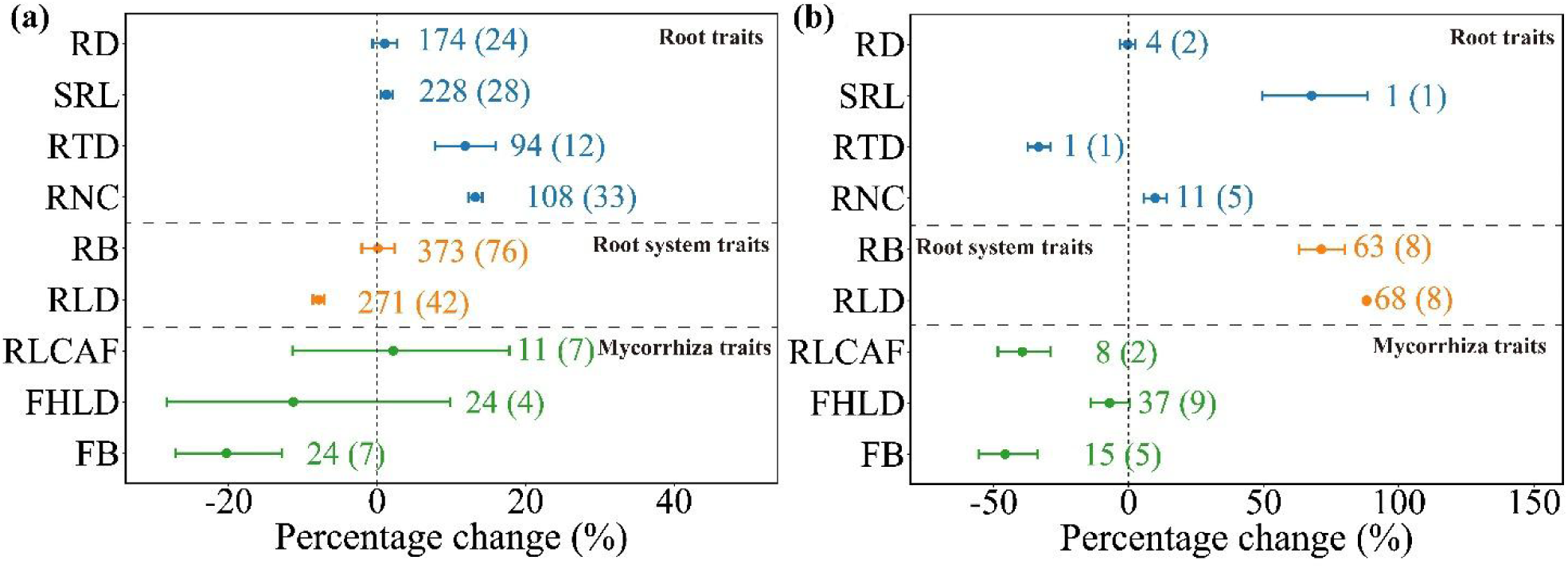
Effect sizes of fine root and mycorrhizal fungal traits in woody (panel a) and herbaceous (panel b) plants in response to nitrogen deposition. Error bars indicate 95% confidence intervals (CIs), with effects considered significant when CIs do not overlap with 0. Numbers within and outside the parentheses represent the sample site and observations for each trait.

Herbaceous plants demonstrated aggressive plasticity: Root tissue density reduction (-33.33%) combined with 67.75% SRL surge and 71.34% biomass accumulation drove 88.14% RLD expansion (Fig. 3b). Enhanced exploitation capacity (+9.79% nitrogen content) co-occurred with rapid mycorrhizal disentanglement - 39.40% colonization loss and 45.73% fungal biomass decline, despite marginal hyphal network changes (-7.05%).

### 3.4 Trait coordination governs nitrogen response syndromes

SRL emerged as pivotal regulator of N response strategies (Fig. 4a-b). Low-SRL plants increased biomass investment (R² = 0.13, *p < 0.001*) and RLD (R² = 0.19, *p < 0.001*), whereas high-SRL strategists reduced root biomass and length. Fine root nitrogen content showed differential associations - decoupled from biomass (*p > 0.1)* but positively correlated with root length density (R² = 0.27, *p < 0.01*, Fig. 4c-d), suggesting nitrogen-enhanced roots prioritize spatial exploration over mass accumulation.

**Figure 4.**
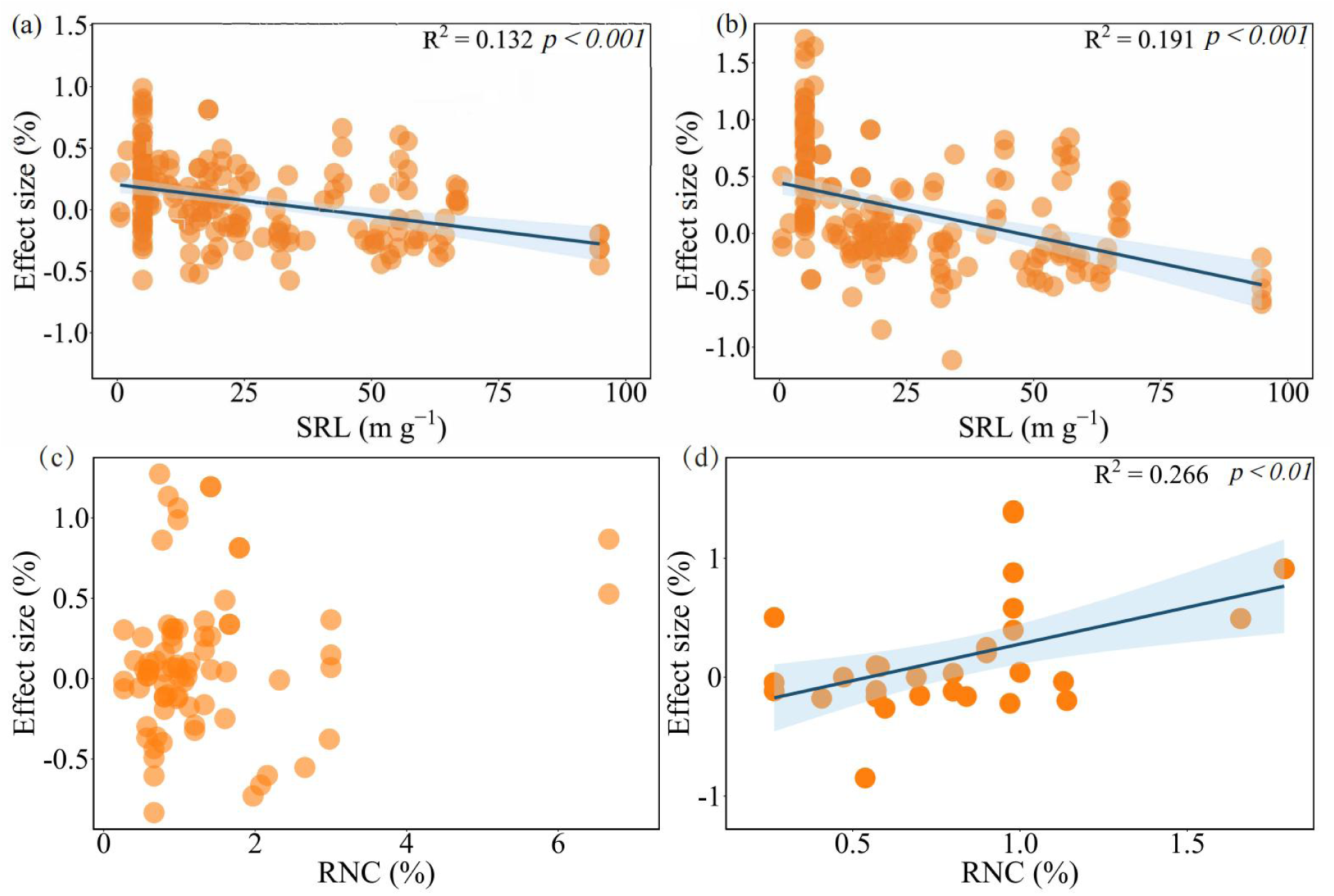
Scatterplot of effect sizes as a function of specific root length (SRL) and root nitrogen content (RNC) for fine root biomass (panel a, panel c) and root length density (panel b, panel d). Trendline for regression relationship are displayed with 95% confidence interval shaded regions, and r-squared and p values above.

### 3.5 Temporal dynamics of nitrogen deposition impacts

N deposition amount positively predicted fine root biomass (R² = 0.015, *p < 0.05*) but negatively impacted fungal biomass (R² = 0.290, *p < 0.05*) (Fig. 5). N deposition duration was significantly negatively correlated with root tissue density (R² = 0.117, *p < 0.001)* (Fig. S4). Additionally, the interaction between N deposition amount and duration negatively impacted fine root biomass and root length density (Table S1).

**Figure 5.**
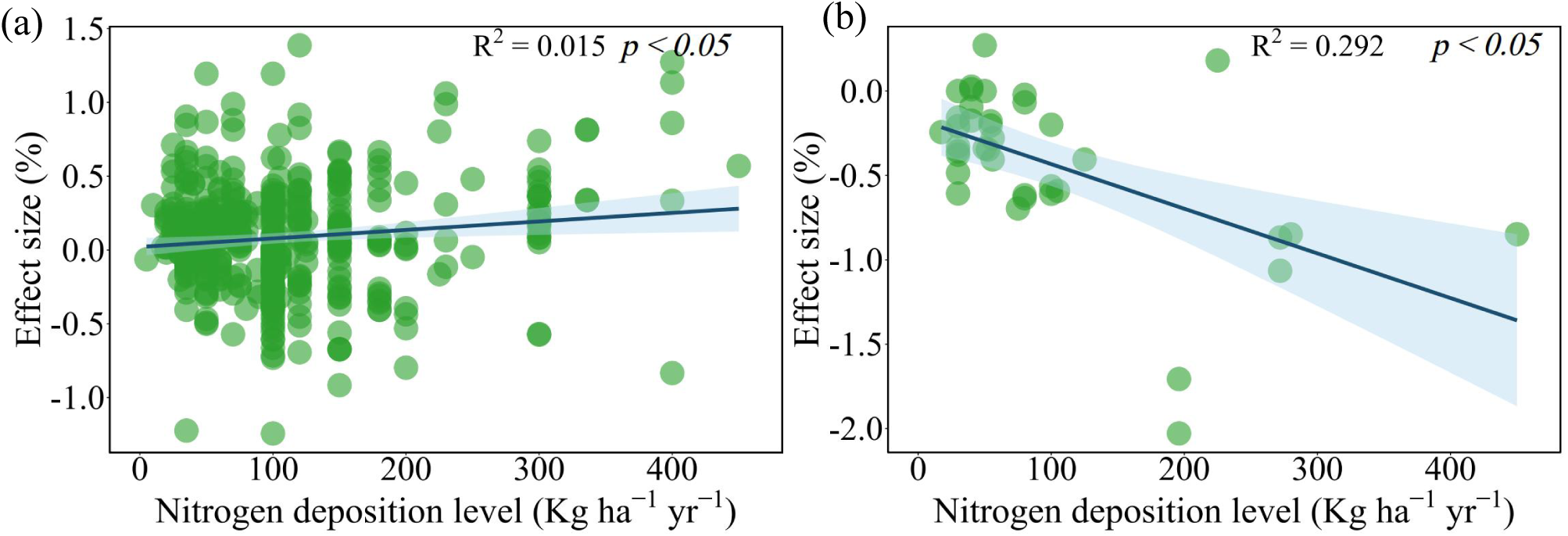
Scatterplot of effect sizes as a function of nitrogen deposition level for fine root biomass (panel a) and mycorrhizal fungal biomass (panel b). Trendline for regression relationship are displayed with 95% confidence interval shaded regions and r-squared and p values above.

## 4. Discussion

### 4.1 Nitrogen deposition reconfigures plant nutrient acquisition paradigms

Contrary to our initial hypothesis (a), nitrogen (N) deposition enhanced both nutrient exploration and exploitation capacities (Fig. 1), a paradox resolved by disentangling growth-form specific responses. Herbaceous species drove this enhancement through radical root architectural remodeling (Fig. 3b), while woody plants exhibited constrained morphological adjustments (-7.87% root length density) (Fig. 3a). This unexpected stability in root morphology suggests that the woody plants’ existing root networks were sufficient to meet the higher nutrient availability without the need for further structural expansion. Instead of allocating energy to grow new roots, the woody plants appeared to enhance nutrient uptake through physiological adjustments. The 13.2% elevation in fine root nitrogen content corresponds with accelerated enzymatic uptake kinetics (Griffiths et al., 2021; Han & Zhu, 2021), effectively supporting 20-35% aboveground biomass gains reported under N enrichment (LeBauer & Treseder, 2008). This metabolic optimization contrasts sharply with herbaceous species’ exploitative strategy: their 88% root length density expansion and 71.34% biomass increase (Fig. 3b) represent extreme phenotypic plasticity to capitalize on ephemeral N pulses. Such rapid belowground restructuring facilitates accelerated reproductive investments, completing life cycles within compressed temporal windows (Salguero-Gómez et al., 2016).

The systematic mycorrhizal disentanglement observed across plant types (-32.8% biomass, -18.73% colonized root length, -7.55% hyphal length) confirms evolutionary cost-benefit optimization in nutrient-rich environments. While mycorrhizal networks provide critical nutrient subsidies in oligotrophic soils (Heijden et al., 2015), their maintenance imposes substantial carbon costs (Han et al., 2021). As N deposition elevates soil nutrient levels, root-autonomous N acquisition becomes energetically favorable, driving the observed fungal biomass reductions.

Divergent responses emerged between mycorrhizal guilds in woody plants (Fig. S2). Arbuscular mycorrhizal (AM) species, predominantly occupying phosphorus-limited tropical ecosystems (Steidinger et al., 2019), countered N-induced P scarcity through 18.7% root length density increases - a compensatory mechanism for impaired fungal P mobilization (Chen et al., 2023). Conversely, ectomycorrhizal (EM) species native to N-limited boreal systems (Du et al., 2020) displayed nitrogen saturation symptoms (Schrijver et al., 2008), reducing root exploration investments as soil N availability surpassed biological demand.

### 4.2 Stoichiometric modulation through phosphorus co-addition

The combined nitrogen and phosphorus (N+P) additions reactivated mycorrhizal mutualisms, evidenced by 41.94% fungal biomass resurgence (Fig. 2), thereby confirming hypothesis (c). This phosphorus-mediated symbiosis revival stems from fundamental stoichiometric constraints: as N deposition elevates plant N:P ratios beyond critical thresholds (Peñuelas et al., 2013), phosphorus emerges as the primary growth-limiting nutrient. Mycorrhizal fungi, particularly efficient in mobilizing immobilized soil phosphorus (Schachtman et al., 1998), regain ecological relevance under these colimitation conditions (Genre et al., 2020). The observed fungal proliferation likely represents strategic resource reallocation - plants reinvest 12-18% of N-induced productivity gains into fungal networks to secure scarce phosphorus (Taiz & Zeiger, 2002).

Notably, N+P co-addition elicited diametrically opposed effects on woody plants’ root architectures compared to N-only treatments (Fig. S3). While solitary N deposition marginally influenced exploration capacity (-7.9%, root length density), N+P synergism drove 25.5% root length density elevation through architectural optimization - reduced tissue density (-7.17%) coupled with enhanced specific root length (+8.99%). This divergence reflects phosphorus’s unique biogeochemical behavior: unlike nitrate’s mass flow-driven mobility (Moreau et al., 2019), phosphate immobilization necessitates active soil exploration. Consequently, woody plants deploy extensive root networks to exploit spatially discrete phosphorus reserves, supporting 25-40% greater aboveground biomass accumulation versus N-only treatments (Jiang et al., 2019).

### 4.3 Root economic space as response predictor

Our findings substantiate hypothesis (d), as plants with lower specific root length (SRL) and greater mycorrhizal dependence showed stronger increases in fine root biomass and root length density under N deposition (Fig. 4a-b). Conversely, plants with higher SRL and autonomous nutrient acquisition strategies reduced their investment in fine root biomass while significantly decreasing root length density. This systematic pattern indicates that N deposition weakens plant-mycorrhizal symbiosis, driving plants to reallocate resources toward developing autonomous root systems to compensate for reduced fungal-mediated nutrient acquisition. Although high-SRL plants typically maintain greater root length density for effective soil exploration (Comas et al., 2012), their limited response under N deposition suggests efficient utilization of existing root networks in nutrient-enriched conditions.

The carbon cycling implications of these responses are significant. Under N deposition, low-SRL plants increase carbon allocation to belowground systems to enhance root nutrient acquisition, while high-SRL plants reduce such investments (Fig. 4a). As fine root biomass constitutes a major component of belowground carbon storage (Jackson et al., 1997), these contrasting strategies validate trait-based approaches for understanding how root functional differences shape ecosystem processes (Díaz et al., 2007) and improve ecological predictions (McCormack et al., 2013). Our findings particularly highlight SRL’s critical role in mediating plant responses to global change, emphasizing the need to integrate fine root traits into ecosystem models.

Fine root nitrogen content further modulated nutrient exploration responses (Fig. 4d). Plants with higher nitrogen content - typically associated with greater photosynthetic demands - expanded root length density to meet elevated nutrient requirements. In contrast, plants with lower nitrogen content showed weaker root length density responses. This pattern parallels the leaf economic spectrum, where fast-growing species gain competitive advantages under N enrichment through increased biomass allocation (Leishman et al., 2010; Ordóñez & Olff, 2013).

Although incomplete trait datasets prevented principal component analysis, significant correlations between root responses, SRL, and nitrogen content allowed approximate positioning within Bergmann et al.’s (2020) root economics framework (Fig. 6). Plants in the lower-right quadrant of this economic space exhibited stronger positive responses to N deposition, while those in the upper-left quadrant showed weaker or negative responses, demonstrating the predictive value of root trait positioning for assessing N deposition impacts.

**Figure 6.**
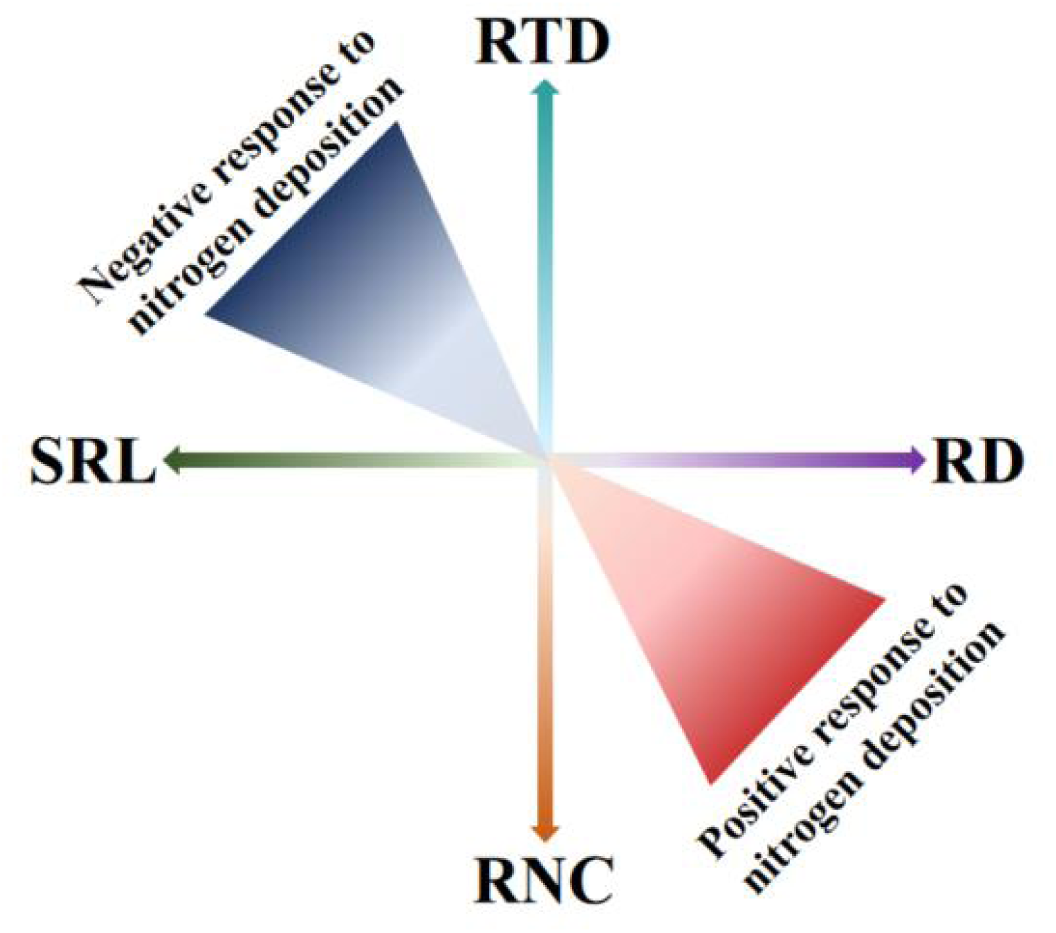
Schematic diagram illustrating the position of fine root traits in the root economics space and their directional responses to nitrogen deposition. The arrows indicate specific root traits: SRL (specific root length), RTD (root tissue density), RD (root diameter), and RNC (root nitrogen content). Colors range from light to dark, reflecting increasing effect sizes of root system trait responses to nitrogen deposition. Positive and negative responses are indicated along the respective axes.

### 4.4 Carbon-nutrient feedback dynamics

Our dose-response analysis revealed a fundamental trade-off between autotrophic and symbiotic nutrient acquisition pathways under nitrogen (N) deposition. Progressive N enrichment drove systematic increases in fine root biomass alongside concomitant declines in mycorrhizal fungal biomass (Fig. 5), corroborating earlier reports of N-induced microbial suppression (Zhang et al., 2018). However, our observed root proliferation contrasts with null effects reported in recent studies (Zhao et al., 2022; Gao et al., 2023), suggesting context-dependent allocation strategies. This divergence likely originates from differential plant investment patterns - as N saturation reduces fungal partnership efficacy (Deng et al., 2017), vegetation compensates through root system expansion to maintain nutrient uptake capacity. The inverse biomass trajectories ultimately reflect a mechanistic shift from fungal-mediated to root-autonomous foraging in N-replete soils.

### 4.5 Limitations and future directions

While our meta-analysis synthesizes global patterns of belowground responses to N deposition, three methodological and conceptual limitations warrant caution in result interpretation. First, observed discrepancies between theoretical predictions of specific root length (SRL) dynamics and empirical data challenge current root trait coordination frameworks. According to the allometric relationship (SRL = 4/[π×D²×RTD]), increasing root tissue density (RTD) under constant diameter should inherently reduce SRL. However, our dataset from woody plants revealed paradoxical SRL elevation (+1.5%) concurrent with RTD increases (+11.8%), suggesting unaccounted mechanisms such as adaptive root branching plasticity or methodological biases in diameter measurement protocols. While our stringent data filtering prioritized high-sensitivity responses, it may have inadvertently excluded critical phenotypic variability, constraining the generalizability of trait coordination patterns. Second, pronounced geographical bias in dataset distribution limits ecological extrapolation. Over 70% of studies originated from temperate ecosystems (Fig. S1), while tropical and boreal biomes remained critically underrepresented. Future syntheses should prioritize filling these geographical knowledge gaps to resolve context-dependent nutrient strategy thresholds. Finally, our isolated focus on N deposition precludes assessment of synergistic or antagonistic interactions with concurrent global change drivers. Emerging evidence suggests elevated CO₂ and warming may counteract observed mycorrhizal suppression by altering plant-fungal carbon trade dynamics (Du et al., 2020), while drought stress could amplify root proliferation responses through hydraulic niche partitioning (Freschet et al., 2021). Incorporating these interactive axes into experimental frameworks will be essential for predicting ecosystem trajectories under multifactorial global change scenarios. Addressing these limitations through standardized trait measurement protocols, expanded geographical sampling, and multifactorial manipulative studies will strengthen mechanistic understanding of anthropogenic N impacts on terrestrial nutrient cycling.

## 5. Conclusion

This study elucidates the mechanistic reorganization of plant nutrient acquisition under nitrogen (N) deposition, revealing a systemic transition from mycorrhizal symbiosis to root-autonomous strategies. Our meta-analysis demonstrates that N enrichment markedly enhances root-mediated nutrient exploration (+84.6% root length density increase) and exploitation (+13.2% nitrogen content elevation) while suppressing mycorrhizal dependence (-32.8% biomass, -18.73% colonized root length, -7.55% hyphal length), marking a fundamental shift in belowground resource allocation (Fig. 7a-b). Divergent lifeform adaptations emerge: woody plants optimize uptake efficiency via nitrogen-enriched roots (+13.2%) despite constrained exploration capacity (-7.9% root length density), whereas herbaceous species exhibit synergistic enhancement of both exploration (+88% root length density) and exploitation (+9.8% nitrogen content and +71.3% root biomass) traits (Fig. 7a-b). Notably, phosphorus co-addition counteracts mycorrhizal suppression (+41.9% biomass), underscoring stoichiometric regulation of plant-fungal interactions in N+P environments. Fine root traits critically predict response magnitudes across species (Fig. 7c-d). High specific root length (SRL) genotypes with lower nitrogen content demonstrate negative responses, contrasting with low-SRL, nitrogen-rich phenotypes showing robust positive plasticity. These findings collectively establish the root economic space as a predictive framework for global change adaptation, wherein trait-mediated thresholds govern carbon-nutrient tradeoffs. In conclusion, this study provides valuable insights into belowground ecological processes under global change and offers a solid theoretical foundation for predicting changes in ecosystem functions.

**Figure 7.**
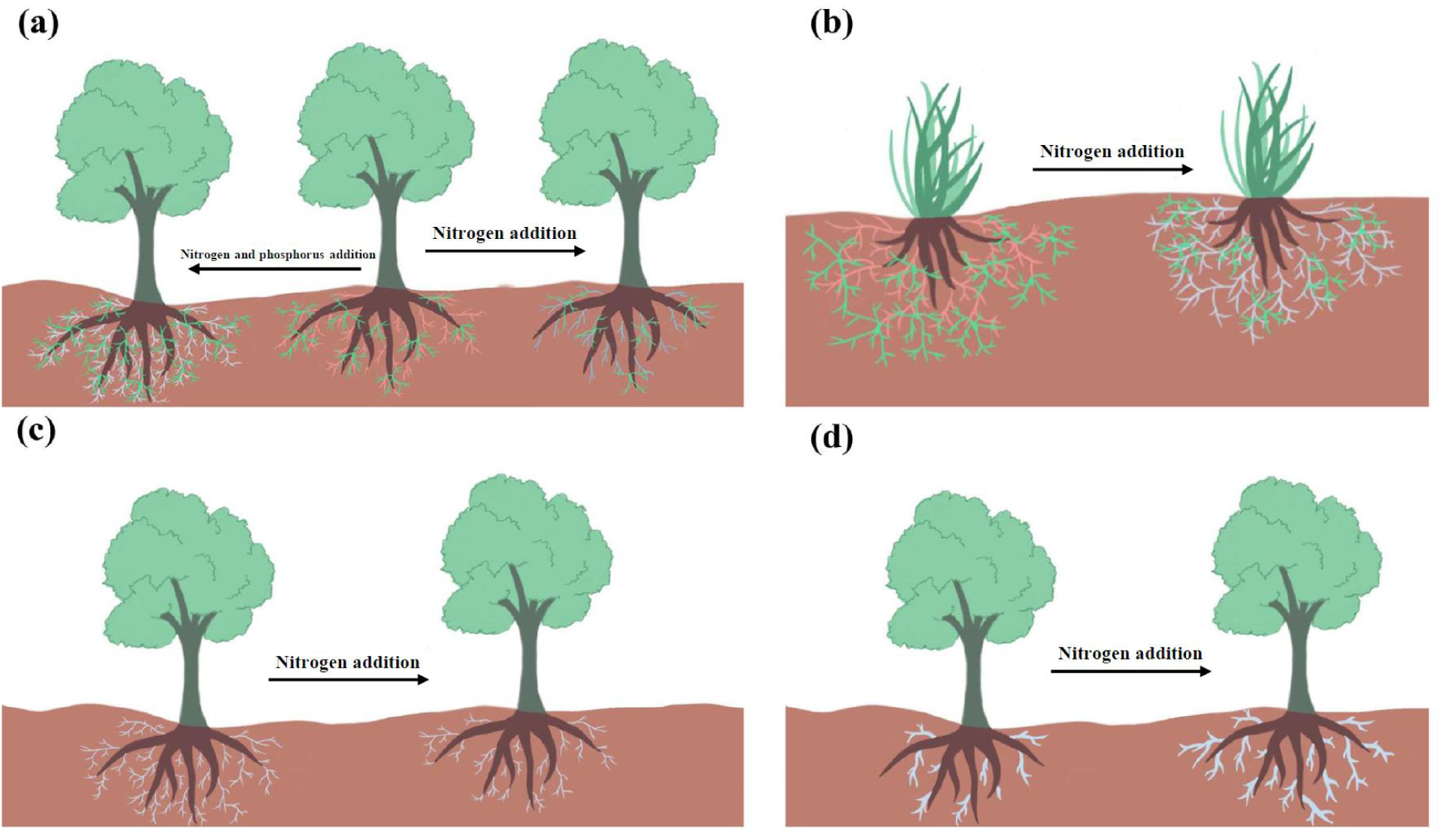
Conceptual diagram of the effects of nitrogen deposition on plant nutrient acquisition strategies. (a) Nitrogen deposition reduces root exploration capacity and dependence on mycorrhizal fungi (depicted by reduced green hyphae) while enhancing root exploitation capacity, as indicated by increased root nitrogen content (root color transitioning from brown to light blue). Nitrogen and phosphorus addition (left panel) enhanced root exploration (longer root lengths) and exploitation (higher root nitrogen content) capacity, with greater reliance on mycorrhizal fungi. (b) In herbaceous plants, nitrogen deposition increases both root exploration and exploitation capacity while reducing dependence on mycorrhizal fungi. (c) Nitrogen deposition causes plants with higher specific root length to reduce root length and biomass. (d) Plants with lower specific root length exhibit increases in root length and biomass under nitrogen deposition.

## Acknowledgements

This study was supported by the Natural Science Foundation of Xiamen, China (grant nos. 3502Z202471075), Xiamen Institute of Technology High level Talent Project (grant no. YKJ24012R).

## Author contributions

H.G. conceived the idea and performed the data analysis. H.G. and Y.Z. wrote the manuscript. Y.Z. Y.H., L.W., and X.Q.C substantially contributed to revisions.

## Data availability

Supplementary data to this article can be found online.

