## Supplementary material for "Nitrogen deposition reshapes plant nutrient acquisition strategies: a meta-analysis within the root economics space": Supplrmrntal materials

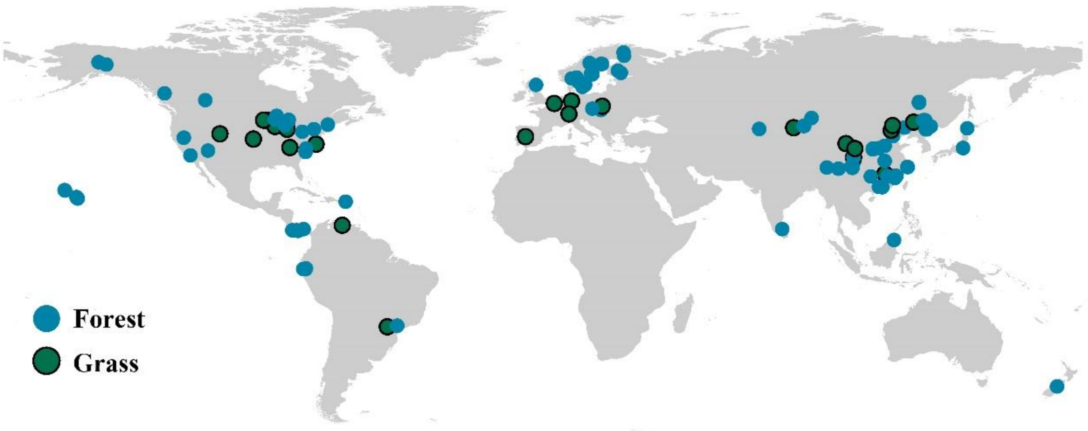


**Figure S1 Global distribution of 153 sites across 135 studies**

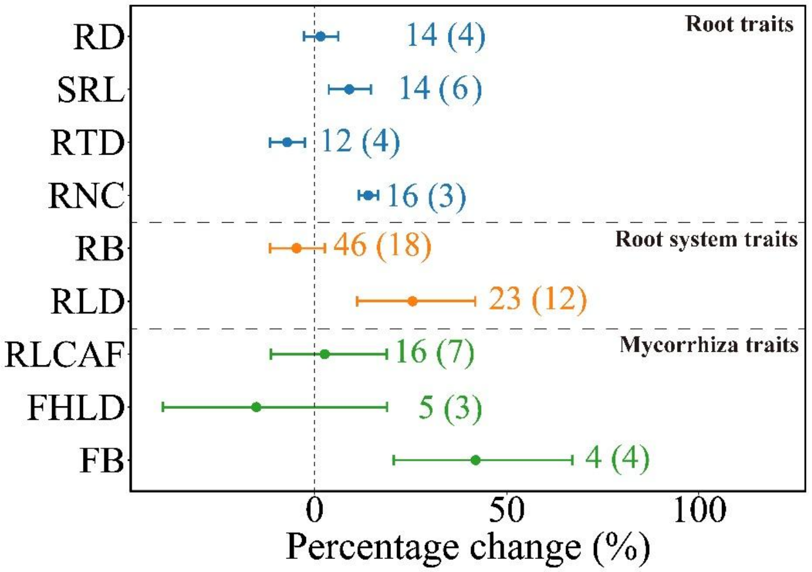


**Figure S2 Effect sizes of nitrogen and phosphorus addition on fine root and mycorrhizal fungal traits for woody plants.** Error bars indicate 95% confidence intervals (CIs), and effects are considered significant when CIs do not overlap with 0. Numbers within and outside the parentheses represent the sample site and observations for each trait.

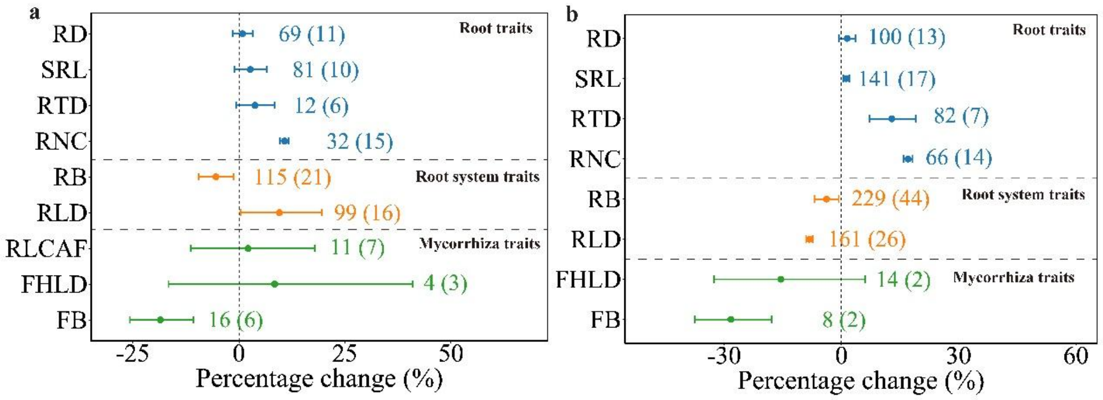


**Figure 3 Effect sizes of nitrogen addition on fine root and mycorrhizal fungal traits in AM (a) and EM (b) woody plants.** Error bars indicate 95% confidence intervals (CIs), with effects considered significant when CIs do not overlap with 0. Numbers within and outside the parentheses represent the sample site and observations for each trait.

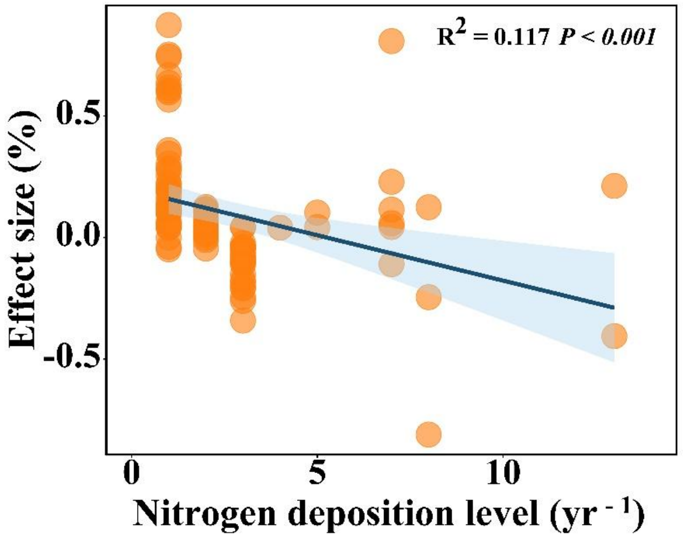


**Figure 4 Scatterplot of effect size of fine root tissue density as a function of nitrogen deposition duration.** Trendline for regression relationship across effect size are displayed with 95% confidence interval (shaded regions) and r-squared and p values above.

.

**Table S1. Significance analysis of nitrogen deposition amount, duration, and their interaction on fine root and mycorrhizal traits based on a mixed-effects model. Bold values indicate significant effects at p < 0.05. The table presents the effect sizes along with corresponding p-values for each trait.**

| | N | D | N*D |
|---|---|---|---|
| Root diameter | 0.001 (*p= 0.43*) | 0.063 (*p= 0.40*) | -0.000 (*p= 0.18*) |
| Specific root length | -0.002 (*p= 0.25*) | 0.033 (*p= 0.56*) | 0.001 (*p= 0.07*) |
| Root tissue density | -0.002 (*p= 0.55*) | -0.904 (***p< 0.001***) | 0.000 (*p= 0.50*) |
| Root nitrogen content | 0.001 (*p= 0.95*) | 0.016 (*p= 0.62*) | 0.000 (*p= 0.07*) |
| Root biomass | 0.002 (***p< 0.05***) | 0.015 (*p= 0.08*) | -0.001 (***p< 0.01***) |
| Root length density | 0.003 (*p= 0.07*) | 0.049 (*p= 0.43*) | -0.001 (***p< 0.05***) |
| Root length colonized by AM fungi | -0.008 (*p= 0.32*) | 0.132 (*p= 0.57*) | -0.001 (*p=0.75*) |
| Fungal hyphal length density | 0.012 (*p= 0.55*) | 0.261 (*p= 0.38*) | -0.003 (*p= 0.36*) |
| Fungal biomass | -0.012 (***p< 0.01***) | 0.007 (*p= 0.90*) | 0.000 (*p= 0.68*) |

A total of 135 studies were included in this meta-analysis for data extraction


1 Ahlström, K., Persson, H. & Börjesson, I. Fertilization in a mature Scots pine (Pinus sylvestris L.) stand—effects on fine roots. *Plant and Soil* **106**, 179-190 (1988).

2 Persson, H. & Ahlström, K. The effects of forest liming on fertilization on fine-root growth. *Water, Air, and Soil Pollution* **54**, 365-375 (1990).

3 BrÆkke, F. H. Root biomass changes after drainage and fertilization of a low-shrub pine bog. *Plant and Soil* **143**, 33-43 (1992).

4 Majdi, H. & Persson, H. Effects of ammonium sulphate application on the chemistry of bulk soil, rhizosphere, fine roots and fine-root distribution in a Picea abies (L.) Karst. stand. *Plant and Soil* **168**, 151-160 (1995).

5 Weber, E. P. & Day, F. P. The effect of nitrogen fertilization on the phenology of roots in a barrier island sand dune community. *Plant and Soil* **182**, 139-148 (1996).

6 Ostertag, R. Effects of nitrogen and phosphorus availability on fine-root dynamics in Hawaiian montane forests. *Ecology* **82**, 485-499 (2001).

7 Majdi, H. Changes in fine root production and longevity in relation to water and nutrient availability in a Norway spruce stand in northern Sweden. *Tree Physiology* **21**, 1057-1061 (2001).

8 Barger, N. N., D'Antonio, C. M., Ghneim, T., Brink, K. & Cuevas, E. Nutrient Limitation to Primary Productivity in a Secondary Savanna in Venezuela1. *Biotropica* **34**, 493-501 (2002).

9 Brunner, I. *et al.* in *Roots: The Dynamic Interface between Plants and the Earth: The 6th Symposium of the International Society of Root Research, Nagoya, Japan.* 253-264 (2001).

10 Phillips, D. L. *et al.* CO 2 and N-fertilization effects on fine-root length, production, and mortality: a 4-year ponderosa pine study. *Oecologia* **148**, 517-525 (2006).

11 Treseder, K. K., Turner, K. M. & Mack, M. C. Mycorrhizal responses to nitrogen fertilization in boreal ecosystems: potential consequences for soil carbon storage. *Global Change Biology* **13**, 78-88 (2007).

12 Berch, S. M., Brockley, R. P., Battigelli, J. & Hagerman, S. Impacts of repeated fertilization on fine roots, mycorrhizas, mesofauna, and soil chemistry under young interior spruce in central British Columbia. *Canadian Journal of Forest Research* **39**, 889-896 (2009).

13 Rosengren-Brinck, U., Majdi, H., Asp, H. & Widell, S. Enzyme activities in isolated root plasma membranes from a stand of Norway spruce in relation to nutrient status and ammonium sulphate application. *New Phytologist* **129**, 537-546 (1995).

14 Mei, L., Gu, J., Zhang, Z. & Wang, Z. Responses of fine root mass, length, production and turnover to soil nitrogen fertilization in Larix gmelinii and Fraxinus mandshurica forests in Northeastern China. *Journal of Forest Research* **15**, 194-201 (2010).

15 Konôpka, B. & Takáčová, E. Effects of liming and NPK-fertilization on the soil and fine roots in a Norway spruce stand, Nízke Tatry Mts. *Ekológia (Bratislava)* **29**, 28-39 (2010).

16 Głąb, T. & Kacorzyk, P. Root distribution and herbage production under different management regimes of mountain grassland. *Soil and Tillage Research* **113**, 99-104 (2011).

17 Rose, L., Coners, H. & Leuschner, C. Effects of fertilization and cutting frequency on the water balance of a temperate grassland. *Ecohydrology* **5**, 64-72 (2012).

18 Teste, F. P., Lieffers, V. J. & Strelkov, S. E. Ectomycorrhizal community responses to intensive forest management: thinning alters impacts of fertilization. *Plant and Soil* **360**, 333-347 (2012).

19 Soares Filho, C. V. *et al.* Root system and root and stem base organic reserves of pasture Tanzania grass fertilizer with nitrogen under grazing. *Semina: Ciências Agrárias, Londrina* **34(5)**, 2415-2426 (2013).

20 Noguchi, K., Nagakura, J. & Kaneko, S. Biomass and morphology of fine roots of sugi (Cryptomeria japonica) after 3 years of nitrogen fertilization. *Frontiers in Plant Science* **4**, 347 (2013).

21 Taylor, B. N. *et al.* Root length, biomass, tissue chemistry and mycorrhizal colonization following 14 years of CO2 enrichment and 6 years of N fertilization in a warm temperate forest. *Tree Physiology* **34**, 955-965 (2014).

22 Wurzburger, N. & Wright, S. J. Fine-root responses to fertilization reveal multiple nutrient limitation in a lowland tropical forest. *Ecology* **96**, 2137-2146 (2015).

23 Chen, G. *et al.* Effect of nitrogen additions on root morphology and chemistry in a subtropical bamboo forest. *Plant and Soil* **412**, 441-451 (2017).

24 Nair, R., Hertel, M., Luo, Y., Moreno, G. & Reichstein, M. Combined effects of altered N: P stoichometry and trees on Mediterranean savanna root dynamics. *Biogeosciences Discussions* **375,** 12-25 (2018)

25 Zheng, Z. & Ma, P. Changes in above and belowground traits of a rhizome clonal plant explain its predominance under nitrogen addition. *Plant and Soil* **432**, 415-424 (2018).

26 Chen, Z. *et al.* Interactive effect of nitrogen addition and throughfall reduction decreases soil aggregate stability through reducing biological binding agents. *Forest Ecology and Management* **445**, 13-19 (2019).

27 Wang, W., Mo, Q., Han, X., Hui, D. & Shen, W. Fine root dynamics responses to nitrogen addition depend on root order, soil layer, and experimental duration in a subtropical forest. *Biology and Fertility of Soils* **55**, 723-736 (2019).

28 Yan, X.L., Jia, L. & Dai, T. Fine root morphology and growth in response to nitrogen addition through drip fertigation in a Populus× euramericana "Guariento" plantation over multiple years. *Annals of Forest Science* **76**, 1-12 (2019).

29 Xiong, D. *et al.* The effects of warming and nitrogen addition on fine root exudation rates in a young Chinese-fir stand. *Forest Ecology and Management* **458**, 117793 (2020).

30 Zhu, H., Zhao, J. & Gong, L. The morphological and chemical properties of fine roots respond to nitrogen addition in a temperate Schrenk's spruce (Picea schrenkiana) forest. *Scientific Reports* **11**, 3839 (2021).

31 Li, W. *et al.* Fine root biomass and morphology in a temperate forest are influenced more by canopy water addition than by canopy nitrogen addition. *Frontiers in Ecology and Evolution* **11**, 1132248 (2023).

32 Wei, G. *et al.* Effects of nitrogen-water interaction on fine root morphology and production in a mixed Pinus koraiensis forest in Changbai Mountains, northeastern China. *Journal of Beijing Forestry University* **38**, 29-35 (2016).

33 He, R.T. *et al.* Effects of combined nitrogen and phosphorus addition on fine root traits of young Machilus pauhoi forest. *The Journal of Applied Ecology* **33**, 337-343 (2022).

34 Peng, Y., Chen, G., Chen, G., Liang, Z. & Tu, L. Effects of simulated nitrogen deposition on soil respiration in a secondary evergreen broad-leaved forest on Wawushan Mountain. *Chinese Journal of Applied and Environmental Biology* **21**, 733-739 (2015).

35 Hou, X. Effect of nitrogen application on soil nitrogen content, root growth and nitrogen absorption of meadow in Wugong mountain. *Pratacultural Science* **35**, 1343-1351 (2018).

36 Gang, H., Xueyong, Z., Yangui, S., Yingxin, H. & Jianyuan, C. Responses of fine root growth of Caragana microphylla shrub plantation to irrigation and nitrogen addition in a semi arid region of northern China. *Journal of Beijing Forestry University* **31**, 73-77 (2009).

37 Kou, L., Guo, D., Yang, H., Gao, W. & Li, S. Growth, morphological traits and mycorrhizal colonization of fine roots respond differently to nitrogen addition in a slash pine plantation in subtropical China. *Plant and Soil* **391**, 207-218 (2015).

38 Liu, Q. *et al.* Belowground responses of Picea asperata seedlings to warming and nitrogen fertilization in the eastern Tibetan Plateau. *Ecological Research* **26**, 637-648 (2011).

39 Noguchi, K. *et al.* Fine-root dynamics in sugi (Cryptomeria japonica) under manipulated soil nitrogen conditions. *Plant and Soil* **364**, 159-169 (2013).

40 Treseder, K. K. & Vitousek, P. M. Effects of soil nutrient availability on investment in acquisition of N and P in

Hawaiian rain forests. *Ecology* **82**, 946-954 (2001).

41 Wang, G., Fahey, T. J., Xue, S. & Liu, F. Root morphology and architecture respond to N addition in Pinus tabuliformis, west China. *Oecologia* **171**, 583-590 (2013).

42 Yin, H. *et al.* Enhanced root exudation stimulates soil nitrogen transformations in a subalpine coniferous forest under experimental warming. *Global Change Biology* **19**, 2158-2167 (2013).

43 Yan, G. *et al.* Nitrogen deposition and decreased precipitation altered nutrient foraging strategies of three temperate trees by affecting root and mycorrhizal traits. *Catena* **181**, 104094 (2019).

44 Fan, Y. *et al.* Responses of soil phosphorus fractions after nitrogen addition in a subtropical forest ecosystem: Insights from decreased Fe and Al oxides and increased plant roots. *Geoderma* **337**, 246-255 (2019).

45 Kårén, O. & Nylund, J.-E. Effects of ammonium sulphate on the community structure and biomass of ectomycorrhizal fungi in a Norway spruce stand in southwestern Sweden. *Canadian Journal of Botany* **75**, 1628-1642 (1997).

46 Nilsson, L. O. & Wallander, H. Production of external mycelium by ectomycorrhizal fungi in a Norway spruce forest was reduced in response to nitrogen fertilization. *New Phytologist* **158**, 409-416 (2003).

47 Johnson, N. C., Rowland, D. L., Corkidi, L., Egerton-Warburton, L. M. & Allen, E. B. Nitrogen enrichment alters mycorrhizal allocation at five mesic to semiarid grasslands. *Ecology* **84**, 1895-1908 (2003).

48 Diepen, L. T. A. V., Lilleskov, E. A., Pregitzer, K. S. & Miller, R. M. Decline of arbuscular mycorrhizal fungi in northern hardwood forests exposed to chronic nitrogen additions. *New Phytologist* **176**, 175-183 (2010).

49 Garcia, M. O., Ovasapyan, T., Greas, M. & Treseder, K. K. Mycorrhizal dynamics under elevated CO 2 and nitrogen fertilization in a warm temperate forest. *Plant and Soil* **303**, 301-310 (2008).

50 Denef, K., Roobroeck, D., Wadu, M. C. M., Lootens, P. & Boeckx, P. Microbial community composition and rhizodeposit-carbon assimilation in differently managed temperate grassland soils. *Soil Biology and Biochemistry* **41**, 144-153 (2009).

51 Fan, Y. *et al.* Decreased soil organic P fraction associated with ectomycorrhizal fungal activity to meet increased P demand under N application in a subtropical forest ecosystem. *Biology and Fertility of Soils* **54**, 149-161 (2018).

52 Jach-Smith, L. C. & Jackson, R. D. N addition undermines N supplied by arbuscular mycorrhizal fungi to native perennial grasses. *Soil Biology and Biochemistry* **116**, 148-157 (2018).

53 Sheldrake, M. *et al.* Responses of arbuscular mycorrhizal fungi to long-term inorganic and organic nutrient addition in a lowland tropical forest. *The ISME Journal* **12**, 2433-2445 (2018).

54 Bradley, K., Drijber, R. A. & Knops, J. Increased N availability in grassland soils modifies their microbial communities and decreases the abundance of arbuscular mycorrhizal fungi. *Soil Biology and Biochemistry* **38**, 1583-1595 (2006).

55 Camenzind, T. *et al.* Opposing effects of nitrogen versus phosphorus additions on mycorrhizal fungal abundance along an elevational gradient in tropical montane forests. *Soil Biology and Biochemistry* **94**, 37-47 (2016).

56 Chen, Y.L. *et al.* Six-year fertilization modifies the biodiversity of arbuscular mycorrhizal fungi in a temperate steppe in Inner Mongolia. *Soil Biology and Biochemistry* **69**, 371-381 (2014).

57 Li, L. *et al.* Different responses of absorptive roots and arbuscular mycorrhizal fungi to fertilization provide diverse nutrient acquisition strategies in Chinese fir. *Forest Ecology and Management* **433**, 64-72 (2019).

58 Staddon, P. L., Jakobsen, I. & Blum, H. Nitrogen input mediates the effect of free-air CO2 enrichment on mycorrhizal fungal abundance. *Global Change Biology* **10**, 1678-1688 (2004).

59 Treseder, K. K. & Allen, M. F. Direct nitrogen and phosphorus limitation of arbuscular mycorrhizal fungi: a model and field test. *New phytologist* **155**, 507-515 (2002).

60 Wang, X. *et al.* NP fertilization did not reduce AMF abundance or diversity but alter AMF composition in an alpine grassland infested by a root hemiparasitic plant. *Plant Diversity* **40**, 117-126 (2018).

61 Wilson, G. W., Rice, C. W., Rillig, M. C., Springer, A. & Hartnett, D. C. Soil aggregation and carbon sequestration are tightly correlated with the abundance of arbuscular mycorrhizal fungi: results from long-term field experiments. *Ecology Letters* **12**, 452-461 (2009).

62 Zhang, T., Yang, X., Guo, R. & Guo, J. Response of AM fungi spore population to elevated temperature and nitrogen addition and their influence on the plant community composition and productivity. *Scientific Reports* **6**, 24749 (2016).

63 Zheng, Y. *et al.* Plant identity exerts stronger effect than fertilization on soil arbuscular mycorrhizal fungi in a sown pasture. *Microbial Ecology* **72**, 647-658 (2016).

64 Zheng, Y. *et al.* Differential responses of arbuscular mycorrhizal fungi to nitrogen addition in a near pristine Tibetan alpine meadow. *FEMS Microbiology Ecology* **89**, 594-605 (2014).

65 Alexander, I. & Fairley, R. Effects of N fertilisation on populations of fine roots and mycorrhizas in spruce humus. *Tree Root Systems and Their Mycorrhizas*, 49-53 (1983).

66 Haynes, B. E. & Gower, S. T. Belowground carbon allocation in unfertilized and fertilized red pine plantations in northern Wisconsin. *Tree Physiology* **15**, 317-325 (1995).

67 Clemensson-Lindell, A. & Persson, H. The effects of nitrogen addition and removal on Norway spruce fine-root vitality and distribution in three catchment areas at Gårdsjön. *Forest Ecology and Management* **71**, 123-131 (1995).

68 Helmisaari, H.S. & Hallbäcken, L. Fine-root biomass and necromass in limed and fertilized Norway spruce (Picea abies (L.) Karst.) stands. *Forest Ecology and Management* **119**, 99-110 (1999).

69 Herbert, D. A., Fownes, J. H. & Vitousek, P. M. Hurricane damage to a Hawaiian forest: nutrient supply rate affects resistance and resilience. *Ecology* **80**, 908-920 (1999).

70 Maier, C. & Kress, L. Soil CO2 evolution and root respiration in 11 year-old loblolly pine (Pinus taeda) plantations as affected by moisture and nutrient availability. *Canadian Journal of Forest Research* **30**, 347-359 (2000).

71 Palatova, E. Effect of increased nitrogen depositions and drought stress on the development of Scots pine (Pinus sylvestris L.)–II. Root system response. *J. For. Sci* **48**, 237-247 (2002).

72 Burton, A. J., Pregitzer, K. S., Crawford, J. N., Zogg, G. P. & Zak, D. R. Simulated chronic NO3− deposition reduces soil respiration in northern hardwood forests. *Global Change Biology* **10**, 1080-1091 (2004).

73 Coleman, M. D., Friend, A. L. & Kern, C. C. Carbon allocation and nitrogen acquisition in a developing Populus deltoides plantation. *Tree Physiology* **24**, 1347-1357 (2004).

74 Cleveland, C. C. & Townsend, A. R. Nutrient additions to a tropical rain forest drive substantial soil carbon dioxide losses to the atmosphere. *Proceedings of the National Academy of Sciences* **103**, 10316-10321 (2006).

75 Hungate, B. A., Hart, S. C., Selmants, P. C., Boyle, S. I. & Gehring, C. A. Soil responses to management, increased precipitation, and added nitrogen in ponderosa pine forests. *Ecological Applications* **17**, 1352-1365 (2007).

76 Jourdan, C. *et al.* Fine root production and turnover in Brazilian Eucalyptus plantations under contrasting nitrogen fertilization regimes. *Forest Ecology and Management* **256**, 396-404 (2008).

77 Magill, A. H. *et al.* Ecosystem response to 15 years of chronic nitrogen additions at the Harvard Forest LTER, Massachusetts, USA. *Forest Ecology and Management* **196**, 7-28 (2004).

78 Mo, J. *et al.* Nitrogen addition reduces soil respiration in a mature tropical forest in southern China. *Global Change Biology* **14**, 403-412 (2008).

79 Cusack, D. F., Silver, W. L., Torn, M. S. & McDowell, W. H. Effects of nitrogen additions on above-and belowground carbon dynamics in two tropical forests. *Biogeochemistry* **104**, 203-225 (2011).

80 Tu, L.H. *et al.* Short-term simulated nitrogen deposition increases carbon sequestration in a Pleioblastus amarus plantation. *Plant and Soil* **340**, 383-396 (2011).

81 Wang, C. *et al.* Responses of fine roots and soil N availability to short-term nitrogen fertilization in a broad-leaved Korean pine mixed forest in northeastern China. *Plos One* **7**, e31042 (2012).

82 Hasselquist, N. J., Metcalfe, D. B. & Högberg, P. Contrasting effects of low and high nitrogen additions on soil CO 2 flux components and ectomycorrhizal fungal sporocarp production in a boreal forest. *Global Change Biology* **18**, 3596-3605 (2012).

83 Xia, M., Talhelm, A. F. & Pregitzer, K. S. Chronic nitrogen deposition influences the chemical dynamics of leaf litter and fine roots during decomposition. *Soil Biology and Biochemistry* **112**, 24-34 (2017).

84 Muratore, T. *et al.* Response of Root Respiration to Warming and Nitrogen Addition Depends on Tree Species. *Global Change Biology* **30**, e17530 (2024).

85 Zhu, F., Yoh, M., Gilliam, F. S., Lu, X. & Mo, J. Nutrient limitation in three lowland tropical forests in southern China receiving high nitrogen deposition: insights from fine root responses to nutrient additions. *Plos One* **8**, e82661 (2013).

86 Tu, L. *et al.* Nitrogen addition stimulates different components of soil respiration in a subtropical bamboo ecosystem. *Soil Biology and Biochemistry* **58**, 255-264 (2013).

87 Gao, Q., Hasselquist, N. J., Palmroth, S., Zheng, Z. & You, W. Short-term response of soil respiration to nitrogen fertilization in a subtropical evergreen forest. *Soil Biology and Biochemistry* **76**, 297-300 (2014).

88 Wang, Q. *et al.* N and P fertilization reduced soil autotrophic and heterotrophic respiration in a young Cunninghamia lanceolata forest. *Agricultural and Forest Meteorology* **232**, 66-73 (2017).

89 Yokoyama, D., Imai, N. & Kitayama, K. Effects of nitrogen and phosphorus fertilization on the activities of four different classes of fine-root and soil phosphatases in Bornean tropical rain forests. *Plant and Soil* **416**, 463-476 (2017).

90 Xiong, D. *et al.* Interactive effects of warming and nitrogen addition on fine root dynamics of a young subtropical plantation. *Soil Biology and Biochemistry* **123**, 180-189 (2018).

91 Li, X. *et al.* Nitrogen deposition and increased precipitation interact to affect fine root production and biomass in a temperate forest: Implications for carbon cycling. *Science of the Total Environment* **765**, 144497 (2021).

92 Ji, J. *et al.* Effects of simulated nitrogen deposition on root biomass of subtropical Chinese fir saplings. *Acta Ecol. Sin* **40**, 6118-6125 (2020).

93 Sun, Y., Gu, J.-C., Zhuang, H.-F. & Wang, Z.-Q. Effects of ectomycorrhizal colonization and nitrogen fertilization on morphology of root tips in a Larix gmelinii plantation in northeastern China. *Ecological Research* **25**, 295-302 (2010).

94 Brenner, R. E., Boone, R. D. & Ruess, R. W. Nitrogen additions to pristine, high-latitude, forest ecosystems: consequences for soil nitrogen transformations and retention in mid and late succession. *Biogeochemistry* **72**, 257-282 (2005).

95 Burton, A. J., Jarvey, J. C., Jarvi, M. P., Zak, D. R. & Pregitzer, K. S. Chronic N deposition alters root respiration-tissue N relationship in northern hardwood forests. *Global Change Biology* **18**, 258-266 (2012).

96 Fan, H. *et al.* Nitrogen deposition promotes ecosystem carbon accumulation by reducing soil carbon emission in a subtropical forest. *Plant and Soil* **379**, 361-371 (2014).

97 Kou, L. *et al.* Simulated nitrogen deposition affects stoichiometry of multiple elements in resource-acquiring plant organs in a seasonally dry subtropical forest. *Science of the Total Environment* **624**, 611-620 (2018).

98 Jia, S. *et al.* N fertilization affects on soil respiration, microbial biomass and root respiration in Larix gmelinii and Fraxinus mandshurica plantations in China. *Plant and Soil* **333**, 325-336 (2010).

99 Davis, M. R., Allen, R. B. & Clinton, P. W. The influence of N addition on nutrient content, leaf carbon isotope ratio, and productivity in a Nothofagus forest during stand development. *Canadian Journal of Forest Research* **34**, 2037-2048 (2004).

100 Hong, Z. et al. Response of fine root morphology and anatomical structure of Betula platyphylla and Populus davidiana natural secondary forest to nitrogen deposition in Changbai Mountains. *Acta Ecologica*

*Sinica* **40**, 608-620 (2020).
101 Ataka, M., Sun, L., Nakaji, T., Katayama, A. & Hiura, T. Five-year nitrogen addition affects fine root exudation and its correlation with root respiration in a dominant species, Quercus crispula, of a cool temperate forest, Japan. *Tree Physiology* **40**, 367-376 (2020).
102 Gong, L. & Zhao, J. The response of fine root morphological and physiological traits to added nitrogen in Schrenk's spruce (Picea schrenkiana) of the Tianshan mountains, China. *PeerJ* **7**, e8194 (2019).
103 Beidler, K. V. *et al.* Changes in root architecture under elevated concentrations of CO2 and nitrogen reflect alternate soil exploration strategies. *New Phytologist* **205**, 1153-1163 (2015).
104 Zhang, X. *et al.* Effects of long-term nitrogen addition and decreased precipitation on the fine root morphology and anatomy of the main tree species in a temperate forest. *Forest Ecology and Management* **455**, 117664 (2020).
105 Bu, W.S. *et al.* The contrasting effects of nitrogen and phosphorus fertilizations on the growth of Cunninghamia lanceolata depend on the season in subtropical China. *Forest Ecology and Management* **482**, 118874 (2021).
106 Liu, G. *et al.* Long-term nitrogen addition regulates root nutrient capture and leaf nutrient resorption in Larix gmelinii in a boreal forest. *European Journal of Forest Research* **140**, 763-776 (2021).
107 Wang, D. *et al.* Effects of short-term N addition on plant biomass allocation and C and N pools of the Sibiraea angustata scrub ecosystem. *European Journal of Soil Science* **68**, 212-220 (2017).
108 Yan, G. *et al.* Spatial and temporal effects of nitrogen addition on root morphology and growth in a boreal forest. *Geoderma* **303**, 178-187 (2017).
109 Wang, D. *et al.* Effects of short-term N addition on soil C fluxes in alpine Sibiraea angustata scrub on the eastern margin of the Qinghai-Tibetan Plateau. *Agricultural and Forest Meteorology* **247**, 151-158 (2017).
110 Adamek, M., Corre, M. D. & Hölscher, D. Responses of fine roots to experimental nitrogen addition in a tropical lower montane rain forest, Panama. *Journal of Tropical Ecology* **27**, 73-81 (2011).
111 Yan, G. *et al.* Sequestration of atmospheric CO2 in boreal forest carbon pools in northeastern China: Effects of nitrogen deposition. *Agricultural and Forest Meteorology* **248**, 70-81 (2018).
112 Xiao, Y. *et al.* Soil-nitrogen net mineralization increased after nearly six years of continuous nitrogen additions in a subtropical bamboo ecosystem. *Journal of Forestry Research* **26**, 949-956 (2015).
113 Bowden, R. D. *et al.* Long-term nitrogen addition decreases organic matter decomposition and increases forest soil carbon. *Soil Science Society of America Journal* **83**, S82-S95 (2019).
114 Zhou, S., Xiang, Y., Tie, L., Han, B. & Huang, C. Simulated nitrogen deposition significantly reduces soil respiration in an evergreen broadleaf forest in western China. *Plos One* **13**, e0204661 (2018).
115 Chen, F. *et al.* Effects of N addition and precipitation reduction on soil respiration and its components in a temperate forest. *Agricultural and Forest Meteorology* **271**, 336-345 (2019).
116 Zhang, H., Liu, Y., Zhou, Z. & Zhang, Y. Inorganic nitrogen addition affects soil respiration and belowground organic carbon fraction for a Pinus tabuliformis forest. *Forests* **10**, 369 (2019).
117 Wang, J., Wang, G., Fu, Y., Chen, X. & Song, X. Short-term effects of nitrogen deposition on soil respiration components in two alpine coniferous forests, southeastern Tibetan Plateau. *Journal of Forestry Research* **30**, 1029-1041 (2019).
118 Liu, H. *et al.* Differential response of soil respiration to nitrogen and phosphorus addition in a highly phosphorus-limited subtropical forest, China. *Forest Ecology and Management* **448**, 499-508 (2019).
119 Liang, L. *et al.* Pathways regulating decreased soil respiration with nitrogen addition in a subtropical forest in China. *Water, Air, & Soil Pollution* **230**, 1-10 (2019).
120 Phillips, R. P. & Fahey, T. J. Fertilization effects on fineroot biomass, rhizosphere microbes and respiratory fluxes in hardwood forest soils. *New Phytologist* **176**, 655-664 (2007).
121 Wright, S. J. *et al.* Potassium, phosphorus, or nitrogen limit root allocation, tree growth, or litter production in a

lowland tropical forest. *Ecology* **92**, 1616-1625 (2011).

122 Jiang, X., Cao, L. & Zhang, R. Changes of labile and recalcitrant carbon pools under nitrogen addition in a city lawn soil. *Journal of Soils and Sediments* **14**, 515-524 (2014).

123 Leppälammi-Kujansuu, J. *et al.* Effects of long-term temperature and nutrient manipulation on Norway spruce fine roots and mycelia production. *Plant and Soil* **366**, 287-303 (2013).

124 Helmisaari, H.-S., Saarsalmi, A. & Kukkola, M. Effects of wood ash and nitrogen fertilization on fine root biomass and soil and foliage nutrients in a Norway spruce stand in Finland. *Plant and Soil* **314**, 121-132 (2009).

125 Persson, H. & Ahlström, K. Fine-root response to nitrogen supply in nitrogen manipulated Norway spruce catchment areas. *Forest Ecology and Management* **168**, 29-41 (2002).

126 Wang, C. *et al.* Six-year nitrogen–water interaction shifts the frequency distribution and size inequality of the first-order roots of Fraxinus mandschurica in a mixed mature Pinus koraiensis forest. *Frontiers in Plant Science* **8**, 1691 (2017).

127 Clemensson-Lindell, A. & Persson, H. Fine-root vitality in a Norway spruce stand subjected to various nutrient supplies. *Plant and Soil* **168**, 167-172 (1995).

128 Razaq, M., Salahuddin, Shen, H.-l., Sher, H. & Zhang, P. Influence of biochar and nitrogen on fine root morphology, physiology, and chemistry of Acer mono. *Scientific Reports* **7**, 5367 (2017).

129 Liu, R. *et al.* Plasticity of fine-root functional traits in the litter layer in response to nitrogen addition in a subtropical forest plantation. *Plant and Soil* **415**, 317-330 (2017).

130 Zhang, C. *et al.* Effects of simulated nitrogen deposition on soil respiration components and their temperature sensitivities in a semiarid grassland. *Soil Biology and Biochemistry* **75**, 113-123 (2014).

131 Kou, L. *et al.* Nitrogen addition regulates tradeoff between root capture and foliar resorption of nitrogen and phosphorus in a subtropical pine plantation. *Trees* **31**, 77-91 (2017).

132 Chen, D., Li, J., Lan, Z., Hu, S. & Bai, Y. Soil acidification exerts a greater control on soil respiration than soil nitrogen availability in grasslands subjected to long-term nitrogen enrichment. *Functional Ecology* **30**, 658-669 (2016).

133 Zhang, Q. *et al.* Short-term effects of soil warming and nitrogen addition on the N: P stoichiometry of Cunninghamia lanceolata in subtropical regions. *Plant and Soil* **411**, 395-407 (2017).

134 Iversen, C. M. & Norby, R. J. Nitrogen limitation in a sweetgum plantation: implications for carbon allocation and storage. *Canadian Journal of Forest Research* **38**, 1021-1032 (2008).

135 Xia, M., Talhelm, A. F. & Pregitzer, K. S. Chronic nitrogen deposition influences the chemical dynamics of leaf litter and fine roots during decomposition. *Soil Biology and Biochemistry* **112**, 24-34 (2017).
